# Language extinction triggers the loss of unique medicinal knowledge

**DOI:** 10.1101/2020.12.03.407593

**Authors:** Rodrigo Cámara-Leret, Jordi Bascompte

## Abstract

There are nearly 7,400 languages in the world and over 30% of these will no longer be spoken by the end of the century^1^. So far, however, our understanding of whether language extinction may result in the loss of linguistically-unique knowledge remains limited. Here, we ask to what degree indigenous knowledge of medicinal plants is associated to individual languages and quantify how much indigenous knowledge may vanish as languages and plants go extinct. Focussing on three independent regions that have a high biocultural diversity —North America, northwest Amazonia, and New Guinea—we show that >75% of all 12,495 medicinal plant services are linguistically-unique, i.e., only known to one language. Whereas most plant species associated with linguistically-unique knowledge are not threatened, most languages that report linguistically-unique knowledge are. Our finding of high uniqueness in indigenous knowledge and strong coupling with threatened languages suggests that language loss will be even more critical to the extinction of medicinal knowledge than biodiversity loss.

---

Indigenous people have accumulated a sophisticated knowledge about plants and their services —including knowledge that confers significant health benefits^2^—that is encoded in their languages^3^. Indigenous knowledge, however, is increasingly threatened by language loss and species extinctions^4,5^. On one hand, language disuse is strongly associated to decreases in indigenous knowledge about plants^6^. On the other hand, global change will constrain the geographic ranges of many human-utilized endemic plants and crops^7,8^. Together, language extinction and reductions in useful plant species within the coming century may limit the full potential of nature’s contributions to people and the discovery of unanticipated uses^9^. So far, however, our understanding of the degree to which the loss of indigenous languages may result in the loss of linguistically-unique knowledge and how this risk compares to that posed by ecological extinction has been limited (Fig. 1).

**Fig. 1.**
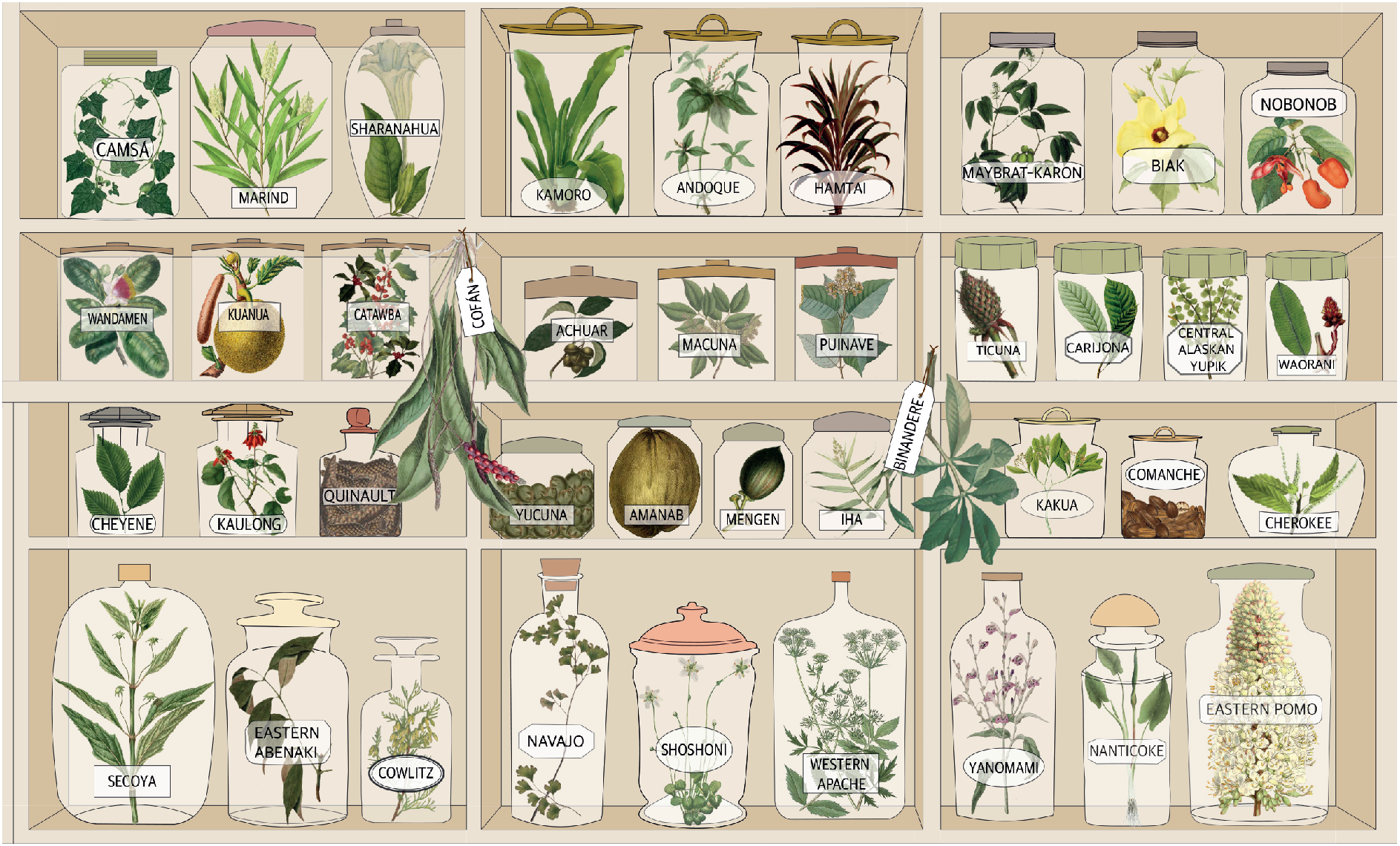
Medicinal plant knowledge and its association to indigenous languages. The figure illustrates a regional pharmacy with remedies (jars with plants) cited by languages (jar labels). In this paper, we assess to what degree the knowledge contained in this pharmacy would be eroded by the extinction of either indigenous languages or plants.

Unravelling the structure of indigenous knowledge about medicinal services has important implications for its resilience^10^. Most indigenous cultures transmit knowledge orally^11^. Therefore, if knowledge about medicines is shared widely amongst indigenous groups that speak different languages, knowledge resilience would be high. That is, even if some indigenous languages go extinct, their medicinal plant knowledge would still be safeguarded in other surviving languages with whom such knowledge is shared. To assess the extent of this, we analyzed three large ethnobotanical datasets for North America^12^, northwest Amazonia^13^, and New Guinea^14^. Together, these data span 3,597 medicinal plant species, and 12,495 plant services associated to 236 indigenous languages (see Methods). We defined a ‘medicinal plant service’ as the combination of a plant species and a medicinal subcategory (e.g., *Ficus insipida* + Digestive System).

Our results show that in all regions, indigenous knowledge about medicinals plants exhibits a strong pattern of linguistic uniqueness, with 73%, 91%, and 84% of the medicinal services in North America, northwest Amazonia, and New Guinea being cited by only one language, respectively (Fig. 2). This finding raises the question of whether unique knowledge is mostly found in languages that are threatened.

**Fig. 2.**
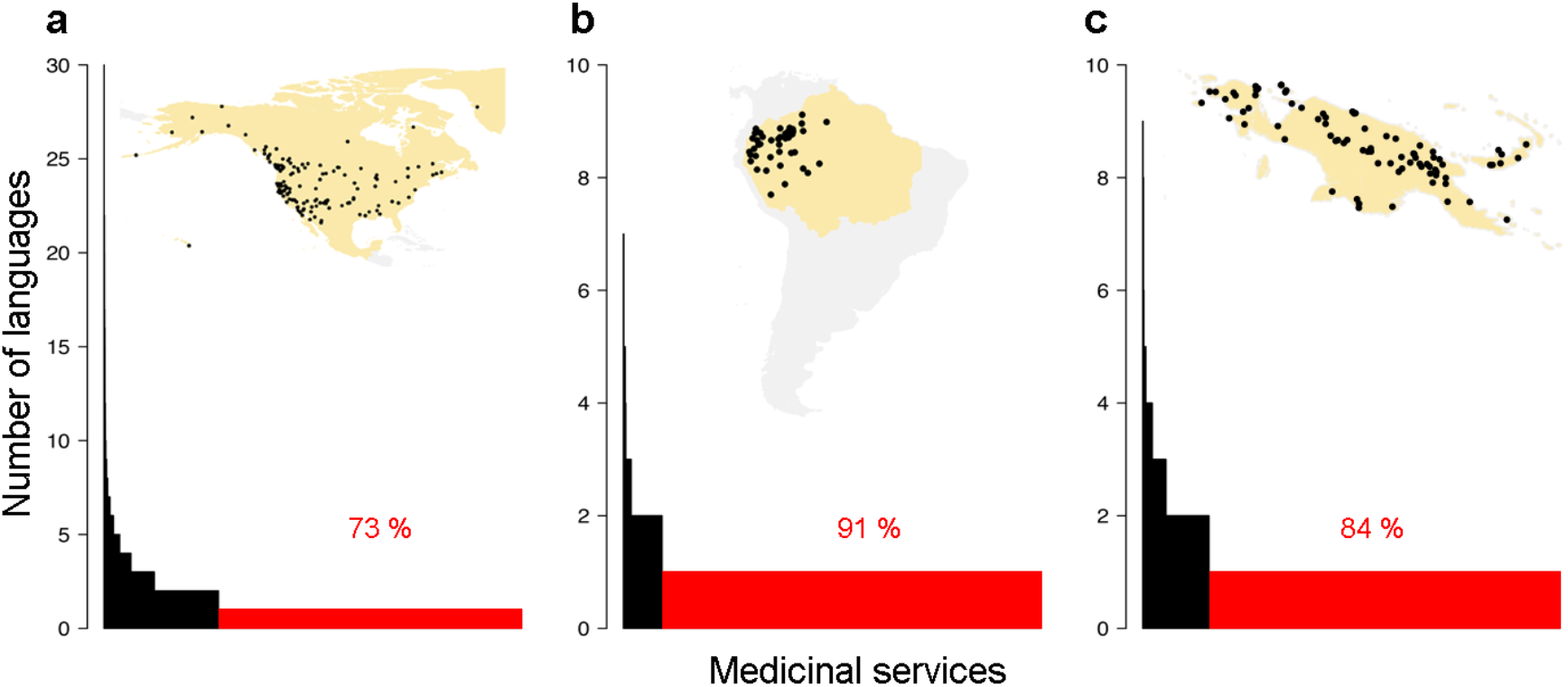
Most medicinal knowledge is unique to a single language. Histograms depict the number of indigenous languages that cite a medicinal service. a, North America; b, northwest Amazonia; c, New Guinea. Red bars show medicinal plant services only known to one language. Dots within the maps indicate the distribution of languages.

Our analysis indicates that threatened languages support 82% and 66% of all unique knowledge in North America and northwest Amazonia, respectively (Supplementary Fig. 1). By contrast, threatened languages account for only 18% of all unique knowledge in New Guinea. This result highlights that the Americas are an indigenous knowledge hotspot (i.e., most medicinal knowledge is linked to threatened languages), and thus a key priority area for future documentation efforts.

Once we have quantified the overall amount of unique knowledge, we next proceed by mapping how it is distributed across the linguistic phylogeny. This will serve to identify whether unique knowledge is uniformly distributed across all linguistic groups, or whether a few linguistic groups deserve more protection than others. First, we built language phylogenies for all the indigenous languages in our sample. Next, we calculated the degree of phylogenetic clustering of unique knowledge using Pagel’s lambda (λ)^15^; values of λ close to 1 indicate strong phylogenetic clustering, whereas values close to 0 indicate data without phylogenetic dependence. We did not find clustering of unique knowledge along the language phylogenies in any of the three regions (Fig. 3, Extended Data Table 1). This indicates that when planning for medicinal knowledge conservation, the entire linguist spectrum —rather than a few “hot” nodes— needs to be considered.

**Fig. 3.**
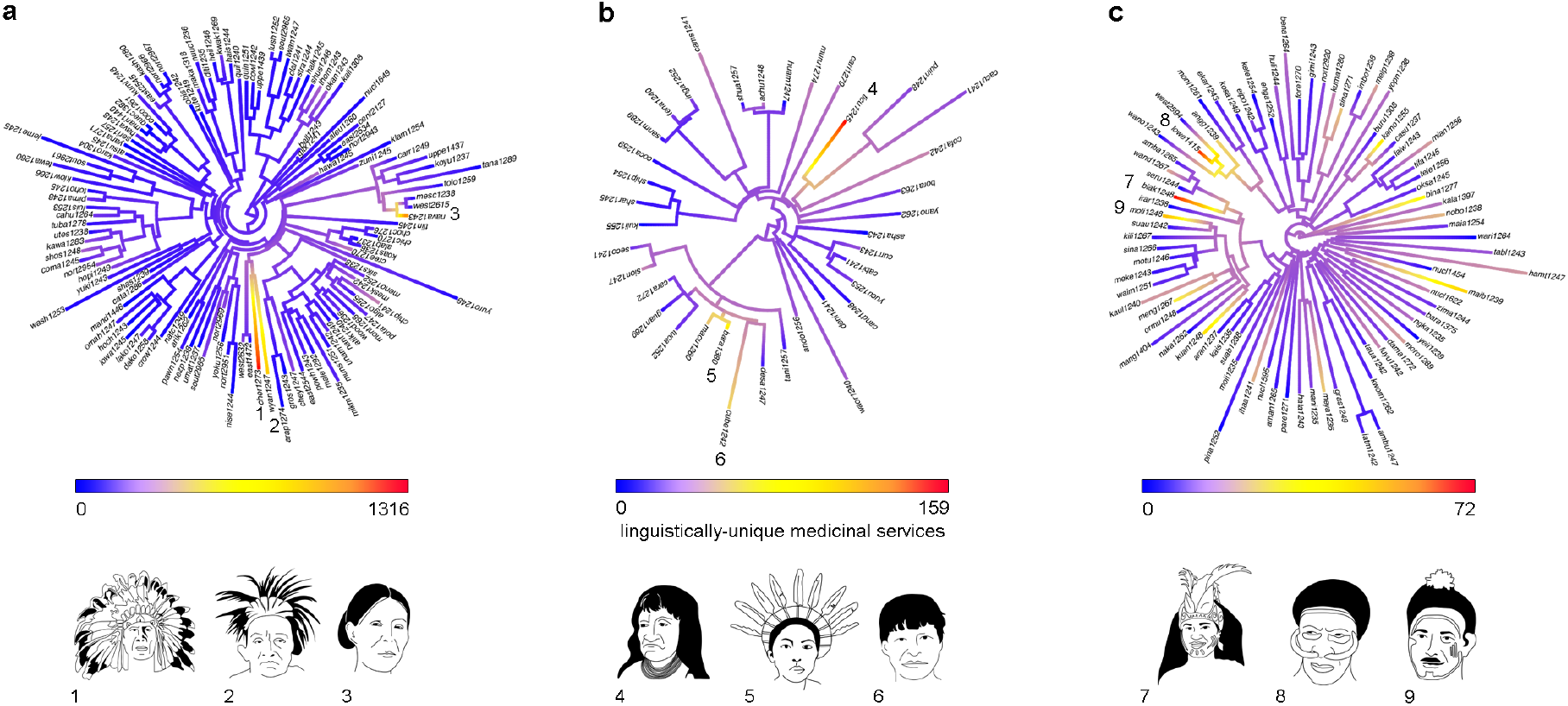
Distribution of unique knowledge across languages. Trees represent language phylogenies of a, North America (*n* = 119 languages); b, northwest Amazonia (*n* = 37 languages); and c, New Guinea (*n* = 80 languages). Illustrations represent indigenous groups whose languages have the highest number of unique medicinal services per region. These languages are indicated by their corresponding numbers in the linguistic trees: 1, Cherokee; 2, Iroquois; 3, Navajo; 4, Tikuna; 5, Barasana; 6, Cubeo; 7, Biak; 8, Lower Grand Valley Dani; 9, Massim. Language names at phylogeny tips are abbreviated following Glottolog codes. For the list of language names and Glottolog codes, see Extended Data Table 2.

So far, we have focused on how unique knowledge is distributed along the cultural dimension. Let us turn now to examine the other component of the indigenous knowledge network, namely the plants. To understand the degree of threat faced by medicinal plants, we queried the IUCN Red List of Threatened species^16^. We found conservation assessments for 22%, 31% and 32% of the medicinal species recorded in North America, northwest Amazonia, and New Guinea, respectively. Of the total medicinal flora with IUCN assessments, 4%, 1%, and 4% were classified as threatened in North America, northwest Amazonia, and New Guinea, respectively (see Methods). To ascertain whether the observed patterns may change as more species are formally assessed, we also obtained conservation predictions from a machine-learning study^17^ (see Methods) which contains assessments for 57%, 25%, and 49% of the medicinal species recorded in North America, northwest Amazonia, and New Guinea, respectively. According to that study, the probability of a medicinal species belonging to a threatened category ranged from 0.0002 to 0.8341 in North America (mean ± SD, 0.156 ± 0.158), 0.149 to 0.822 in northwest Amazonia (mean 0.483 ± 0.119), and 0.063 to 0.679 in New Guinea (mean 0.357 ± 0.141), respectively. In summary, both the IUCN conservation assessments and machine-learning predictions suggest that most medicinal plant species in our sample are not threatened. Finally, we found that less than 1% of all unique knowledge in each region was associated to both threatened languages and threatened plants (Extended Data Table 3). However, there is considerable uncertainty about the potential loss of unique knowledge from the extinction of plants because 61% and 46% of the unique knowledge in North America and northwest Amazonia that is associated to threatened languages belongs to plants that lack plant conservation assessments. IUCN conservation assessments are urgently needed for these plant species.

To assess whether unique knowledge is strongly clustered biologically, we built phylogenies of the medicinal floras of each region, and calculated Pagel’s lambda (Fig. 4). We only found significant clustering of unique knowledge in North America, although values were low (Extended Data Table 1). This relatively weak phylogenetic signal across the three regions suggests that when planning for biocultural conservation, the entire medicinal flora —rather than a few clades— must be considered.

**Fig. 4.**
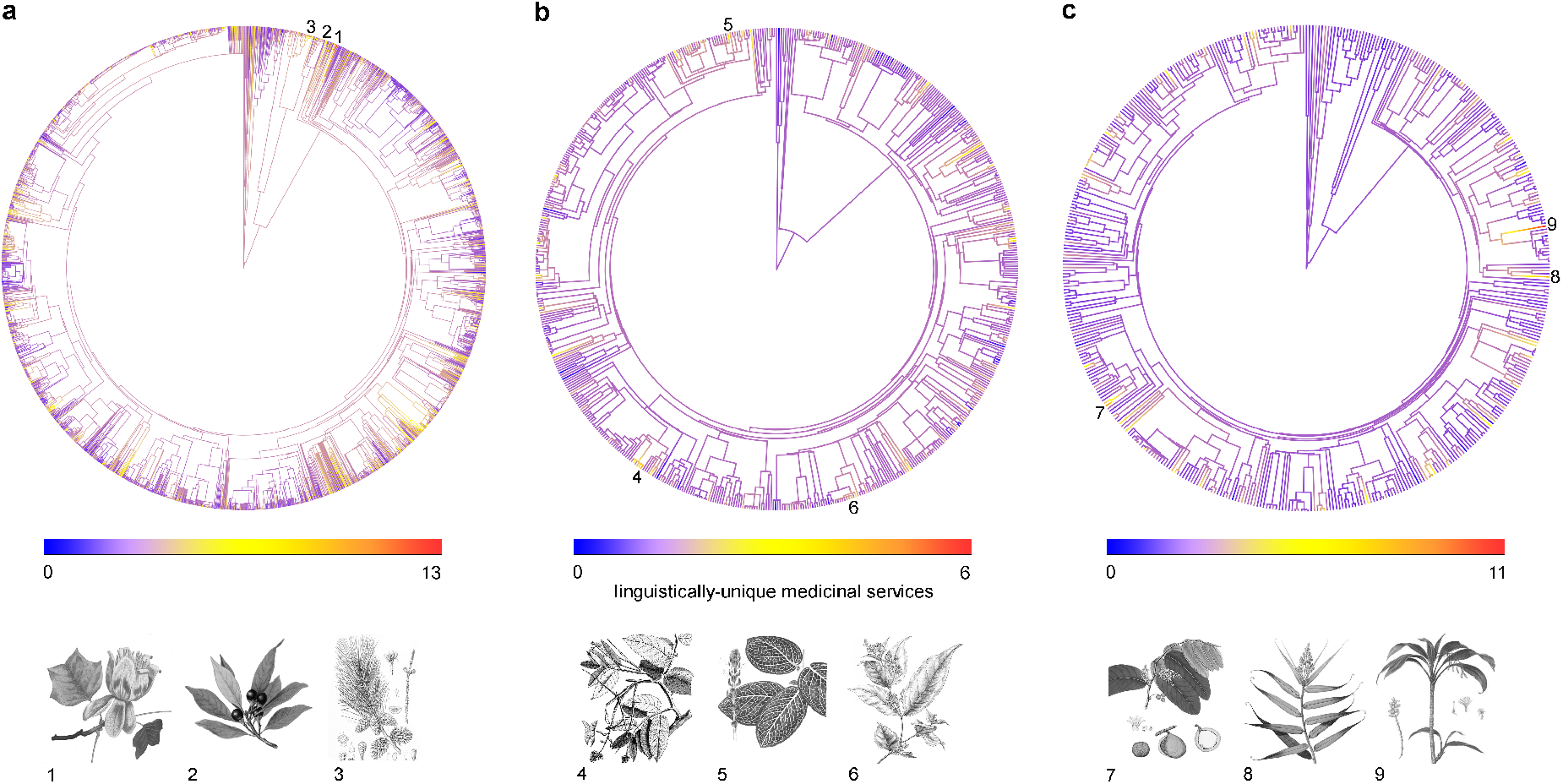
Distribution of unique knowledge across medicinal floras. Trees represent medicinal plant phylogenies of a, North America (n = 2,475 species); b, northwest Amazonia (n = 645 species); and c, New Guinea (n = 477 species). Illustrations and their corresponding numbers show the plant species with more unique medicinal services per region. 1, *Liriodendron tulipifera*; 2, *Persea borbonia*; 3, *Pinus glabra*; 4, *Tachigali paniculata*; 5, *Fittonia albivenis*; 6, *Tetrapterys styloptera*; 7, *Inocarpus fagifer*; 8, *Flagellaria indica*; 9, *Cordyline fruticosa.* All illustrations from http://www.plantillustrations.org belong to the public domain.

Here, we have shown that in North America, northwest Amazonia, and New Guinea, indigenous knowledge of medicinal plant services exhibits a low redundancy across languages that is typical of systems with high information content^18,19^. This low redundancy in medicinal knowledge among languages does not support the notion of high cross-cultural consensus, i.e., that cultures resemble each other in their knowledge, but instead highlights the unique biocultural heritage each culture holds. The invention and diversification of languages involves two opposing forces. On the one hand, sharing facilitates the exchange of information and the spread of valuable ideas that may enhance the fitness within populations. On the other hand, the diversification of languages is the result of innovations, and eventually linguistic barriers may limit information spread. In areas of high linguistic or biological diversity, and/or geographic barriers, the balance between sharing and innovating may tip towards the latter. This may result in the amplification of differences among cultures, as we have shown here for the case of medicinal knowledge.

The United Nations declared 2019 as the year of the world’s Indigenous languages to raise awareness of their endangerment across the world. Our study suggest that each indigenous language brings unique insights that may be complementary to other societies who seek potentially-useful medicinal remedies. Therefore, the predicted extinction of up to 30% of indigenous languages by the end of the 21st century^1^ would substantially compromise humanity’s capacity for medicinal discovery.

## Methods

### Plant Services

We obtained a list of medicinal plant species and services associated to individual indigenous groups from three regions: 1) North America: from the Native American Ethnobotany database^12^— the largest repository of indigenous knowledge for the region; 2) northwest Amazonia: from Richard E. Schultes’s book on the medicinal plants of northwestern Amazonia, which integrates nearly half a century of his field research^13^; and 3) New Guinea: from an ethnobotanical review of 488 references and 854 herbarium specimens^14^.

We classified uses from the three data sources into medicinal subcategories following the classification in the *Economic Botany Data Collection Standard*^20^, with modifications explained by Camara-Leret *et al.*^21^. Medicinal subcategories included Blood and cardiovascular system; Cultural diseases and disorders; Dental health; Digestive system; Endocrine system; General ailments with unspecified symptoms; Infections and infestations; Metabolic system and nutrition; Muscular-skeletal system; Nervous system and mental health; Poisoning; Pregnancy, birth and puerperium; Reproductive system and reproductive health; Respiratory system; Sensory system; Skin and subcutaneous tissue; Urinary system; Veterinary; Not specified; Other medicinal uses. We defined ‘unique knowledge’ as a medicinal service cited exclusively by one indigenous language. By omitting ‘plant parts’ (e.g., bark, leaf, fruit, seed) from our definition of medicinal plant services (i.e.,the combination of plant species and a medicinal subcategory), our categorization is more conservative and underestimates the detection of medicinal knowledge that is restricted to one language.

### Language Phylogenies and Threat

Medicinal services in the literature were associated to 119 indigenous languages in North America, 37 languages in northwest Amazonia, and 80 languages in New Guinea. For each region, we built language trees through phylogenetic inference using machine learning techniques on the word lists of the Automated Similarity Judgement Program (ASJP v.18) and used the Glottolog classification as a constraint tree^22^. To assess the degree of threat faced by languages in our sample, we queried the Ethnologue^23^ which uses the Expanded Graded Intergenerational Disruption Scale (EGIDS) to quantify language threat^24^. For a list of the languages analyzed, see Extended Data Table 2.

### Vascular Plant Phylogenies and Threat

We verified plant species taxonomy using recently published checklists to the vascular plants of the Americas^25^ and New Guinea^26^. Using the list of medicinal plant species in each region, we queried the mega-tree GBOTB.-extended of Smith & Brown^27^ with the *phylo.maker* function of the R package V.PhyloMaker^28^. The phylogenies used in all subsequent analyses comprised 2,475 species in North America, 645 species in northwest Amazonia, and 477 species in New Guinea. To assess the threat faced by medicinal plant species, we queried the conservation assessments published by the IUCN Red List of Threatened species^16^, which classifies species as Data Deficient, Least Concern, Near Threatened, Vulnerable, Endangered, and Critically Endangered, Extinct in the Wild, and Extinct. Following IUCN, species assessed to be Near Threatened, Vulnerable, and Endangered were considered threatened. Because most plant species lack IUCN conservation assessments, we also obtained endangerment probabilities from a recent study that used machine-learning to predict the conservation status of 30,497 plant species^17^.

## Acknowledgments

We thank G. Jäger for providing the language trees, the members of the Bascompte lab for helpful discussions, and I. Cámara Leret for the illustrations in Figs. 1 and 3.

## Author Contributions

R. Cámara-Leret contributed to conceptualization, data collection, data analysis, writing - original draft, writing - review and editing; J. Bascompte contributed to conceptualization, writing - review and editing.

## Supplementary Information

is available for this paper as Extended Data Tables 1-3.

## Correspondence and requests for materials

should be addressed to: Rodrigo Cámara-Leret

---

**Extended Data Table 1.** Phylogenetic clustering (measured using Pagel’s λ) of unique knowledge along the language and plant phylogenies of North America, Northwest Amazonia, and New Guinea. Statistically significant results: ***, P-value < 0.001.

|  | Languages | Plants |
| --- | --- | --- |
| North America | 0.31 | 0.21*** |
| Northwest Amazonia | 6.61e-05 | 6.61e-05 |
| New Guinea | 6.61e-05 | 0.02 |

**Extended Data Table 2.**
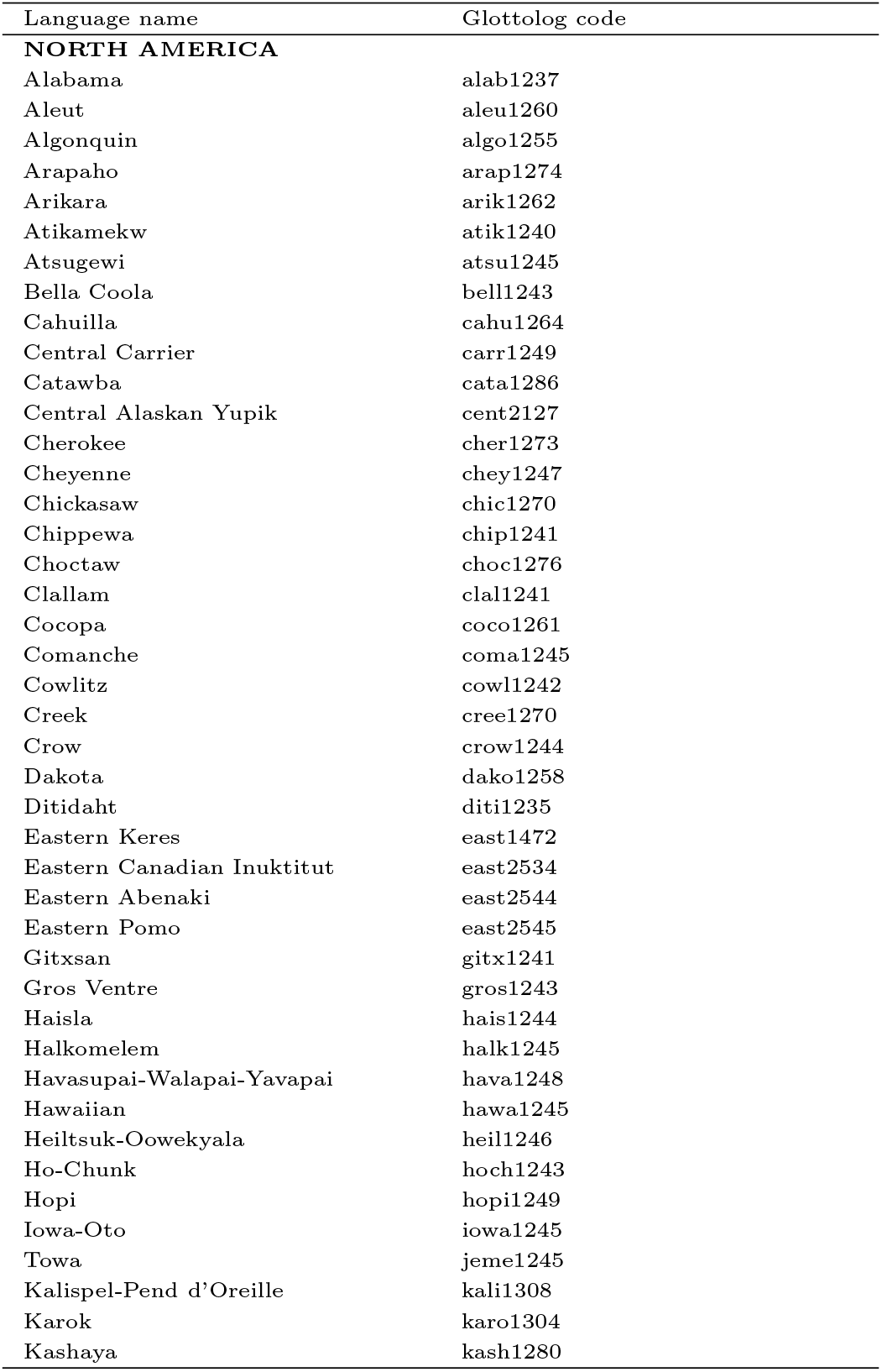
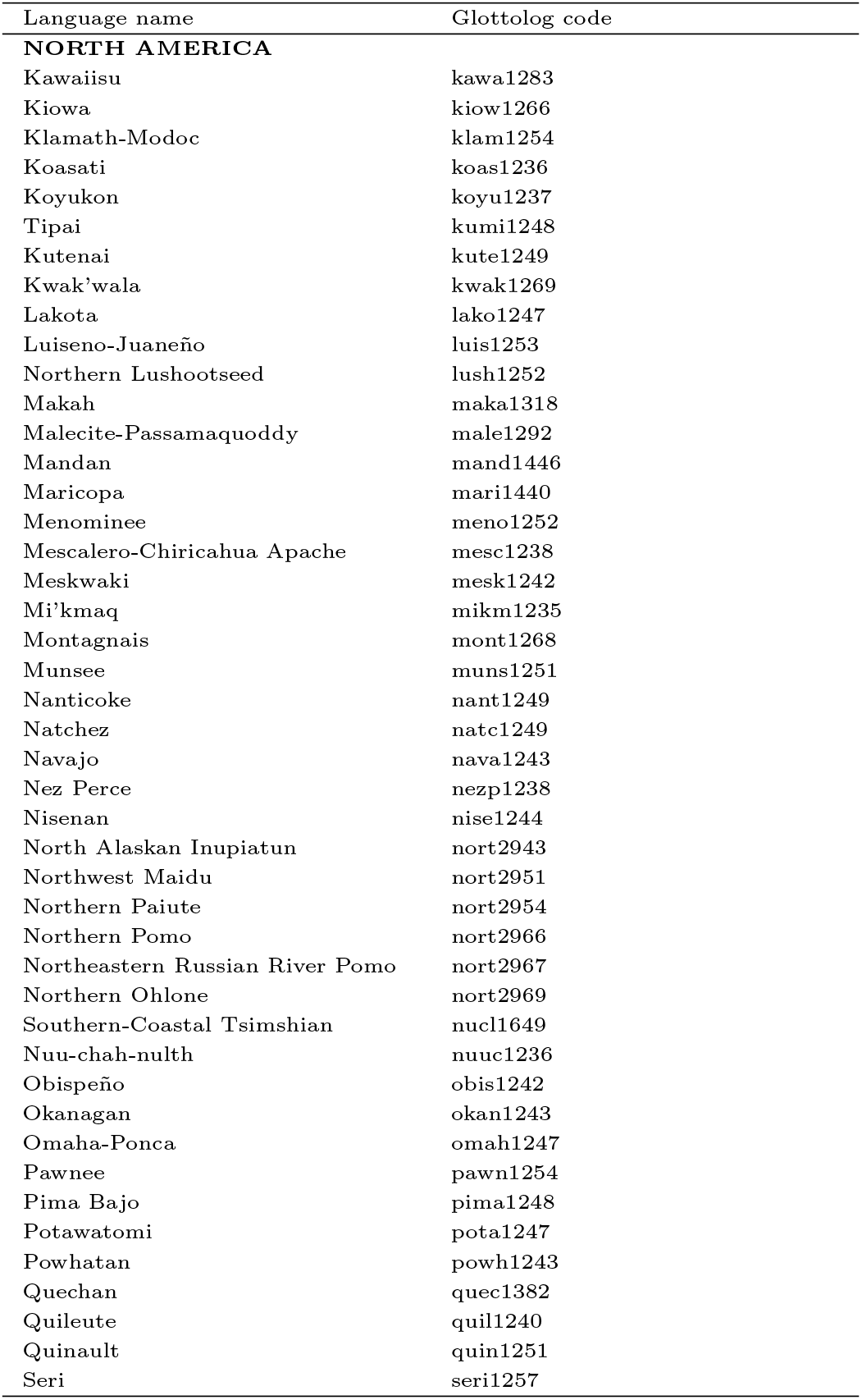
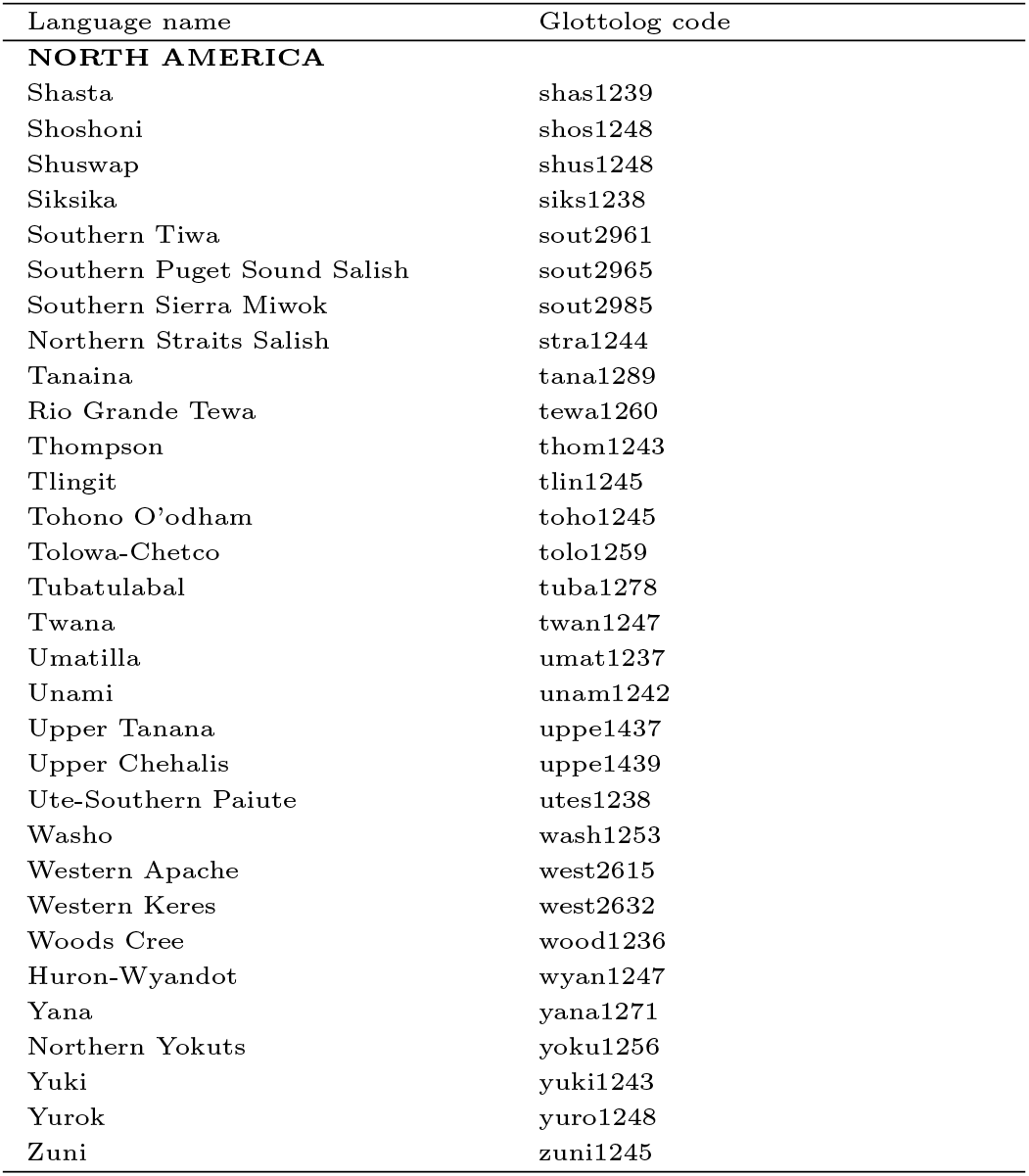
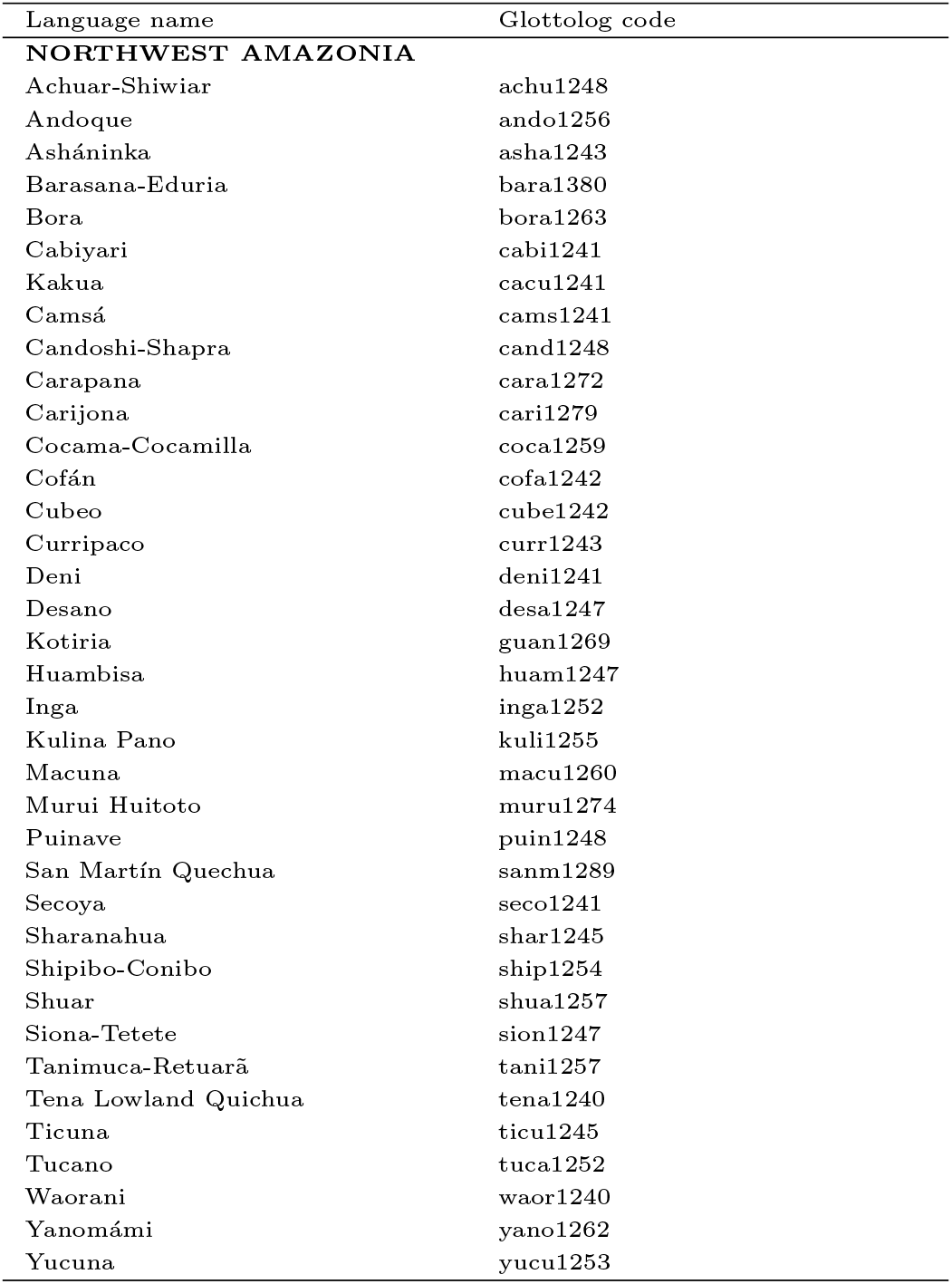
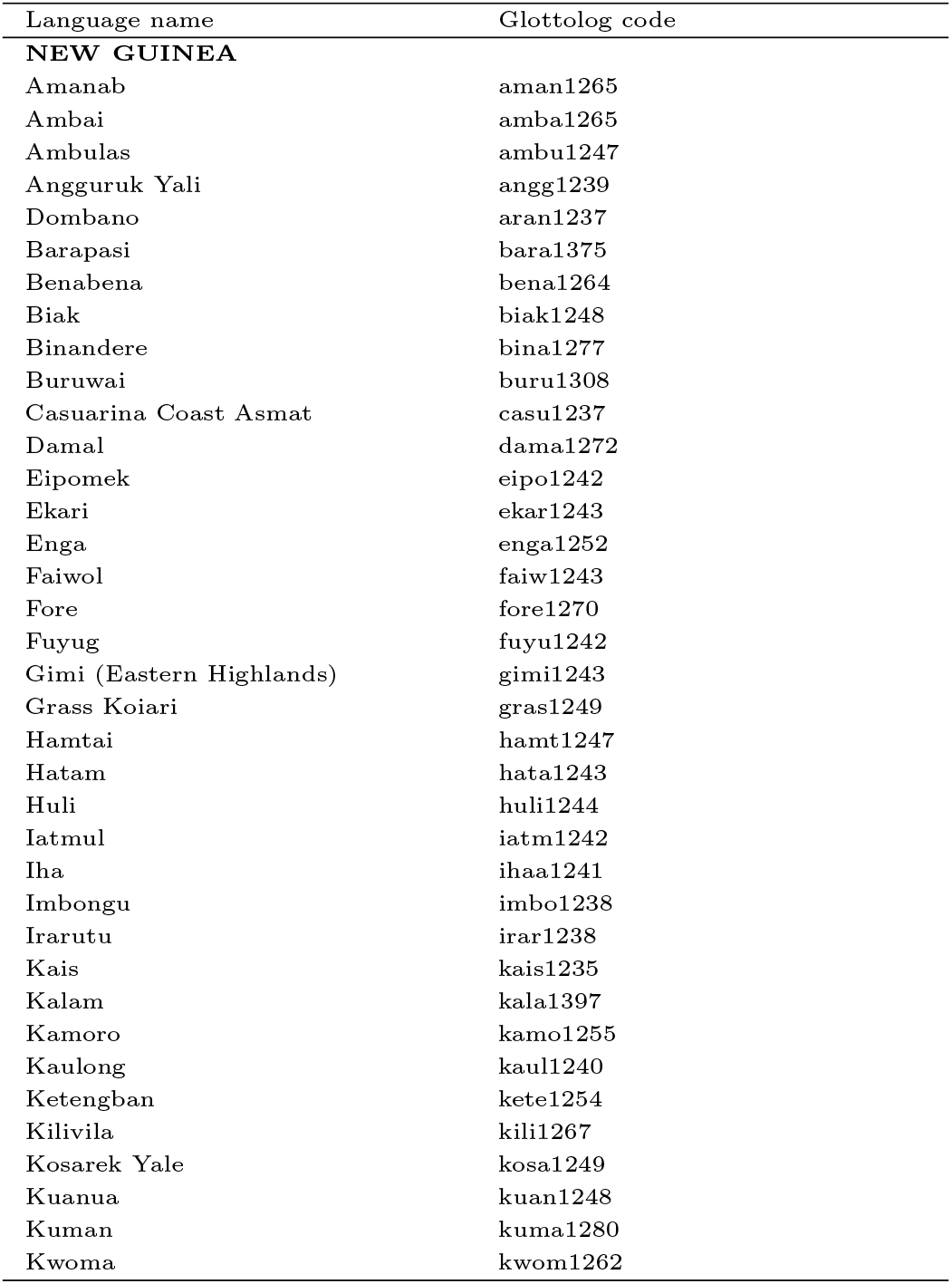
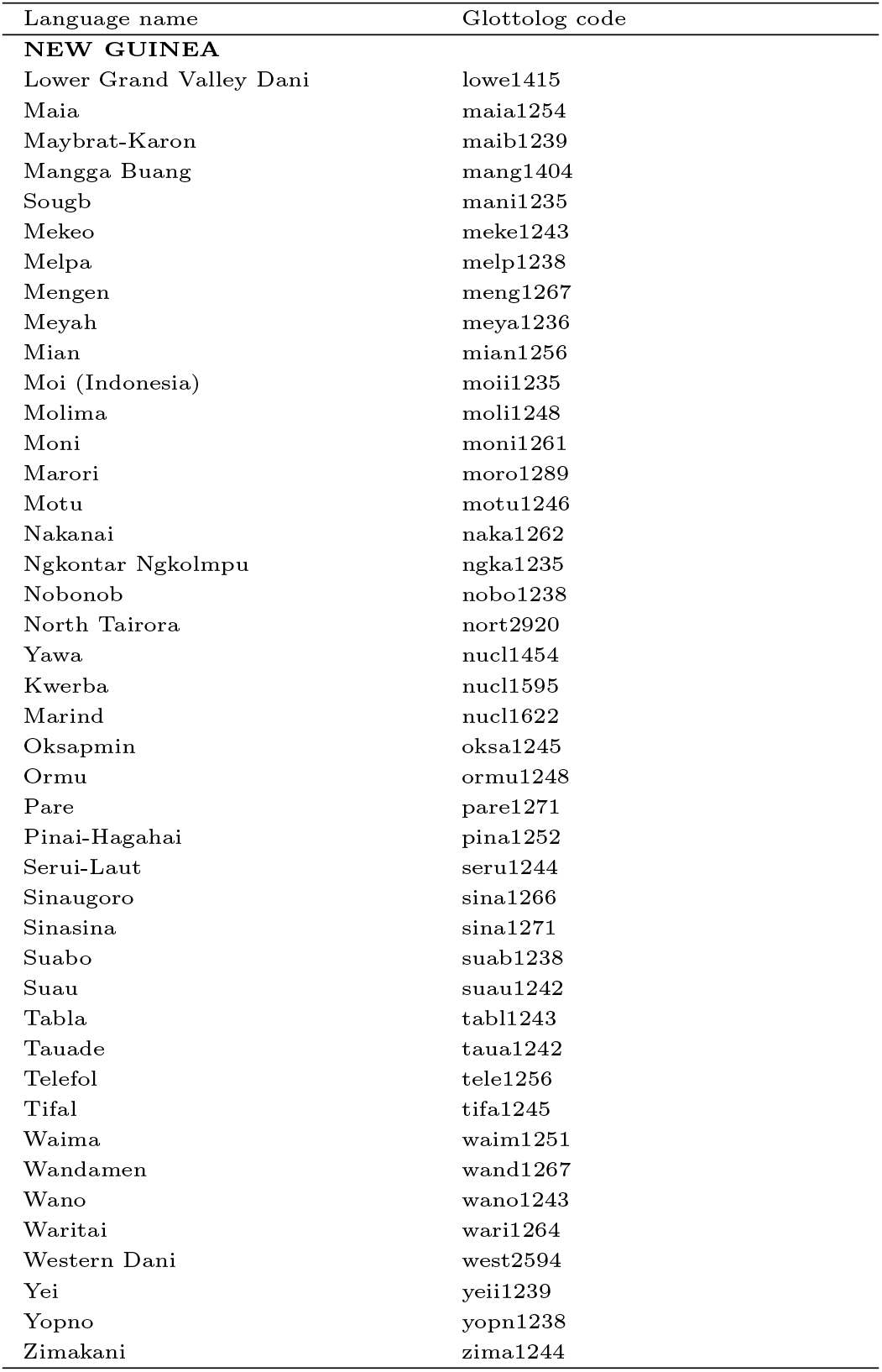
Names and Glottolog codes of the studied languages of North America, northwest Amazonia, and New Guinea.

**Extended Data Table 3.**
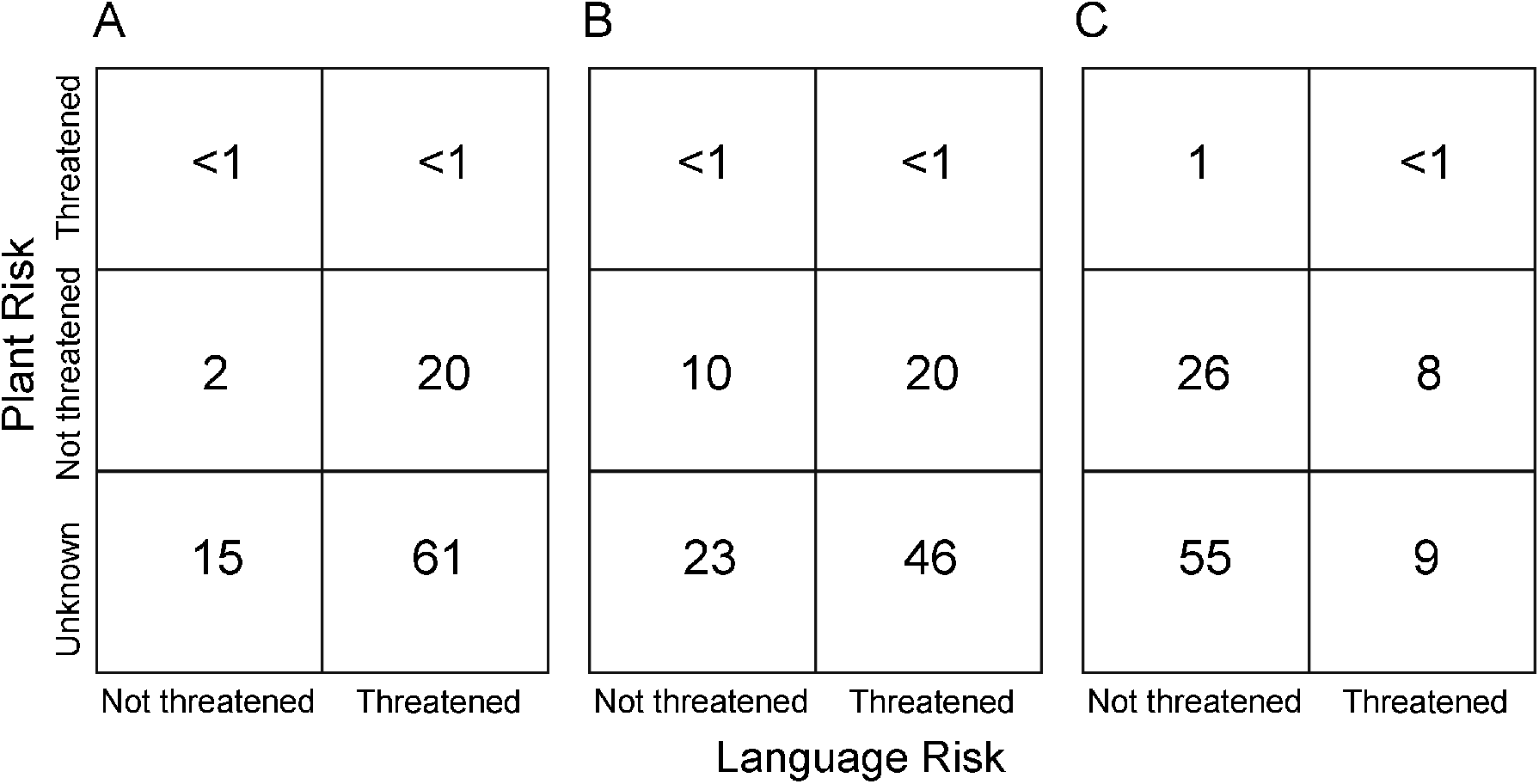
The percentage of unique knowledge associated to threatened and non-threatened languages and plants. a, North America (*n* = 7,565 medicinal services); b, northwest Amazonia (*n* = 773 medicinal services); c, New Guinea (*n* = 873 medicinal services). Language threat follws the classification in the Ethnologue^23^. Plant threat follows the IUCN Red List of Threatened species^16^.

